# Enabling metagenomic surveillance for bacterial tick-borne pathogens using nanopore sequencing with adaptive sampling

**DOI:** 10.1101/2021.08.17.456696

**Authors:** Evan J. Kipp, Laramie L. Lindsey, Benedict S. Khoo, Christopher Faulk, Jonathan D. Oliver, Peter A. Larsen

## Abstract

Technological and computational advancements in the fields of genomics and bioinformatics are providing exciting new opportunities for pathogen discovery and surveillance. In particular, single-molecule nucleotide sequence data originating from Oxford Nanopore Technologies (ONT) sequencing platforms can be bioinformatically leveraged, in real-time, for enhanced biosurveillance of a vast array of zoonoses. The recently released nanopore adaptive sampling (NAS) pipeline facilitates immediate mapping of individual nucleotide molecules (i.e., DNA, cDNA, and RNA) to a given reference as each molecule is sequenced. User-defined thresholds then allow for the retention or rejection of specific molecules, informed by the real-time reference mapping results, as they are physically passing through a given sequencing nanopore. Here, we show how NAS can be used to selectively sequence entire genomes of bacterial tick-borne pathogens circulating in wild populations of the blacklegged tick vector, *Ixodes scapularis*. The NAS method provided a two-fold increase in targeted pathogen sequences, successfully enriching for *Borrelia* (*Borreliella*) *burgdorferi* s.s.; *Borrelia* (*Borrelia*) *miyamotoi*; *Anaplasma phagocytophilum*; and *Ehrlichia muris eauclairensis* genomic DNA within our *I. scapularis* samples. Our results indicate that NAS has strong potential for real-time sequence-based pathogen surveillance.

## 1. Introduction

Nanopore sequencing, based on platforms pioneered by Oxford Nanopore Technologies, Inc. (ONT), has proven to be an increasingly valuable tool for a variety of long-read and metagenomic sequencing applications. In brief, nanopore sequencing is accomplished by passing individual DNA (or RNA) molecules through a membrane-embedded protein nanopore; as a strand of nucleic acid moves through the nanopore, changes in ionic current across the membrane are measured and processed into nucleotide sequence data [1–4]. Importantly, the single-molecule nature of nanopore sequencing permits notable advantages over other next-generation sequencing technologies. In particular the palm-sized ONT MinION and MinION Mk1C sequencing instruments are user-friendly, generate real-time results, and are highly portable, thus enabling their use across a wide variety of lab-based and field-based settings [5–11]. Furthermore, single-molecule nanopore sequencing can yield long-read sequencing data, with individual sequences measuring many thousands of bases in length, without a requirement for clonal library amplification. This aspect of the nanopore sequencing approach helps to limit potential sequencing biases related to complex or repetitive genomic regions where short-read sequencing-by-synthesis methods (e.g., Illumina) typically struggle [12–14]. Although the per-base accuracy of ONT sequencing platforms is less than that of other next-generation platforms, steady advances in sequencing chemistry and basecalling have yielded continued and substantial improvements in reliability [3,4,15].

Due in part to these key advantages—ease of library preparation, the small and portable footprint of ONT sequencing instruments, and the generation of real-time sequencing data— nanopore sequencing is particularly well-suited for the purposes of pathogen detection and biosurveillance [14,16–18]. In contrast to conventional surveillance methods, which rely heavily on nucleic acid amplification (i.e., PCR), pathogen detection by nanopore sequencing can: (1) bypass the requirement that investigators possess *a priori* knowledge of the pathogen(s) being targeted, (2) enable detection of many agents simultaneously, and (3) provide a greater scope of downstream genomic sequence information suitable for pathogen strain typing, functional characterization, and phylogenetics [16,19]. For these reasons, metagenomic and metatranscriptomic approaches using nanopore sequencing offer an innovative and unbiased means for detecting and characterizing a wide variety of bacterial, viral, and eukaryotic pathogens of relevance to both human and animal health [19–21].

Nanopore adaptive sampling (NAS) is a recently developed and novel method that performs selective sequencing of individual DNA, cDNA, or RNA molecules in real-time using ONT sequencing technology [22–24]. NAS operates through an advanced bioinformatic pipeline that compares nucleotides of individual molecules (~200-400 base pairs) against a user-specified reference file every ~0.4 sec, as sequencing is occurring. This dynamic method has a wide number of applications and can selectively enrich targets of interest (e.g., select genes, chromosomes, mitogenomes, RNAs, etc.) and reject unwanted molecules throughout a given sequencing run [23,25]. NAS can be leveraged for real-time pathogen surveillance and discovery using two approaches. First, depletion of host genomic DNA or RNA sequences can be accomplished in real-time by supplying a host reference genome (e.g., human, bat, rodent, primate, mosquito, tick) and rejecting host sequences. The resulting sequence data is enriched for the entire metagenomic community and then mined for *de novo* surveillance of putative pathogens. Second, a reference file containing whole genomes or select genes (e.g., phylogenetically informative genes, antimicrobial resistance genes) of all known pathogens of interest (including both viral and bacterial targets in a single file) can be provided to the NAS pipeline (Figure 1). If present within a biological sample, sequences with a minimum of ~70% sequence similarity to the reference database will be retained [23]. This aspect of NAS distinguishes it from all other sequencing methods, as the process can detect extremely small proportions of non-host DNA/RNA (e.g., low abundance viral and/or bacterial pathogens intermixed with >99% host molecules) [24]. A critical observation with respect to NAS is that the method is not restricted with respect to template length and thus the resulting single-molecule sequences can be thousands of bases long. Thus, the metagenomic data generated by NAS increases confidence in resulting surveillance findings and taxonomic identification as entire viral or bacterial genomes can be recovered. For these reasons, NAS-based surveillance can be used for *de novo* identification of pathogens via unbiased metagenomic/metatranscriptomic enrichment or through the capture of novel strains or species due to sequence similarity shared with suspected agents based on clinical observations (e.g., SARS-CoV-2 shares 82% similarity with SARS) [26].

**Figure 1.**
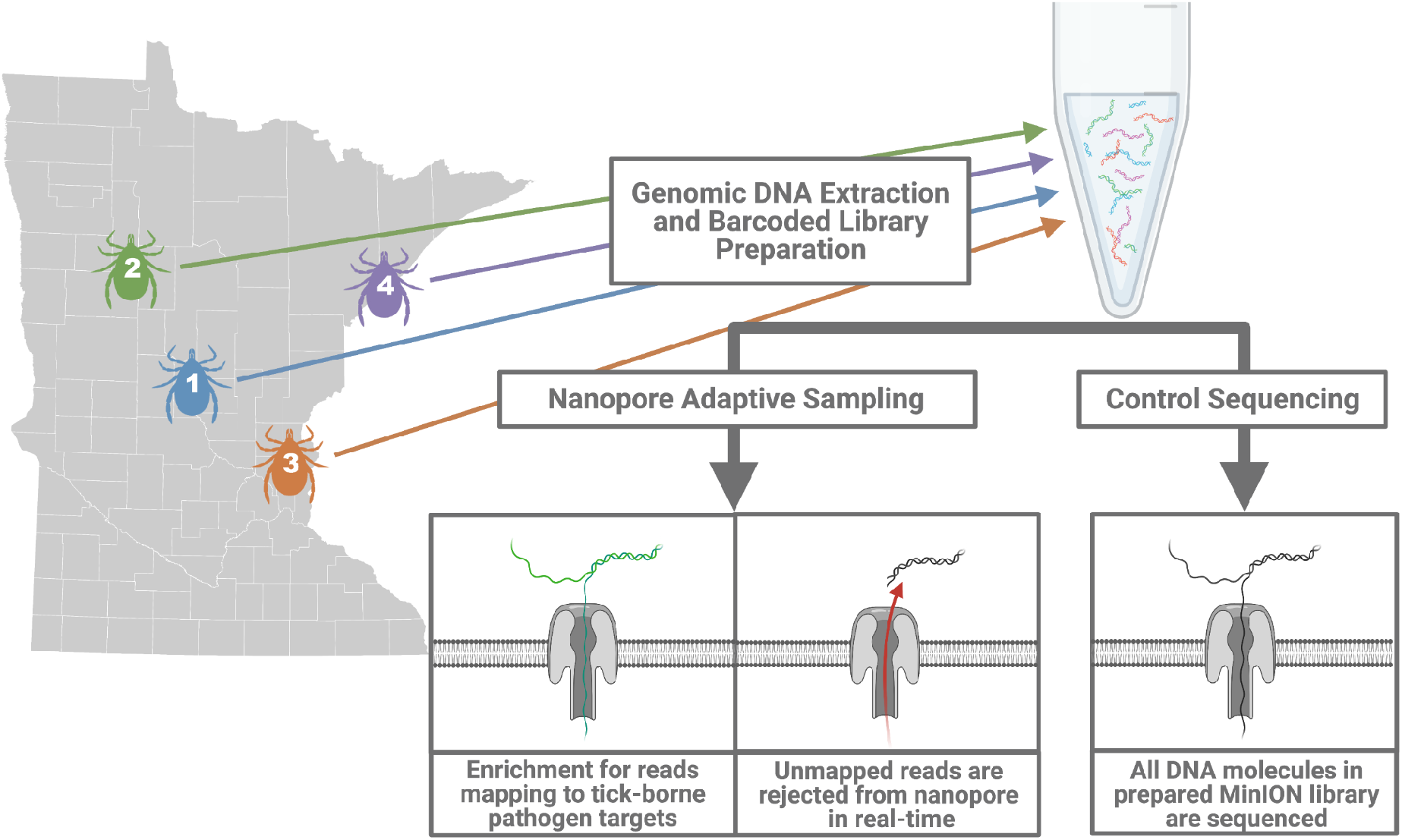
Comparison of nanopore adaptive sampling (NAS) and control sequencing methods. In contrast to control sequencing, in which all molecules of the nanopore library are sequenced, NAS-based surveillance allows the investigator to specify target sequences for either enrichment or depletion. When enriching for whole pathogen genomes, unmapped host-derived reads will be rejected from the sequencing nanopore in real-time to prevent sequencing output on unwanted molecules. Map on the left shows sampling localities in Minnesota, USA where each of the four adult *I. scapularis* specimens was collected for evaluation in this study.

To evaluate the utility of the NAS pipeline for pathogen surveillance, we attempted to detect and characterize tick-borne pathogens (TBPs) vectored by the blacklegged tick, *Ixodes scapularis* (Acari: Ixodidae). Globally, ixodid ticks include some of the most significant disease vectors of importance to both human and animal health, and are capable of transmitting a wide diversity of bacterial, protozoal, and viral pathogens to their vertebrate hosts. As a group, TBPs serve as favorable targets to understand how NAS can be leveraged for sequence-based pathogen surveillance. Recent studies have highlighted the complex bacterial, viral, and eukaryotic metagenomic communities maintained by tick species and populations, which harbor an abundance of both pathogenic and non-pathogenic microorganisms [21,27–31]. Frequently, individual ticks may be co-infected with multiple TBPs, the epidemiological and clinical consequences of which are still poorly understood [27,32,33]. In the United States, *I. scapularis* represents the vector species of greatest public health importance [34]. While *I. scapularis* is best known for its role in the transmission of Lyme disease—caused by *Borrelia (Borreliella) burgdorferi* s.l.—it is increasingly evident that the species is also responsible for transmitting numerous other TBPs. In addition to Lyme disease, at least six other human pathogens are currently recognized to be vectored by *I. scapularis*, including the causative agents of *Borrelia miyamotoi* disease, anaplasmosis, ehrlichiosis, babesiosis, and Powassan virus encephalitis [34–37].

A complicating factor in the control of *I. scapularis*-transmitted diseases has been the dramatic expansion of vector populations over the past decades, concurrent with increasing vector densities and human-tick contact rates throughout many regions of North America [34,38–40]. As a result, the reported incidence of tick-borne diseases in the U.S. over the past two decades has more than doubled [38]. Despite these trends, current methods for tick surveillance are limited in their reliance on conventional PCR and ability to detect only a single or limited range of TBPs. To evaluate the potential of nanopore sequencing—in combination with the NAS pipeline—for tick surveillance, we attempted to detect and characterize TBPs from field-collected *I. scapularis* ticks from Minnesota, USA, a state with a high burden of tick-borne disease [28,41]. Using our approach, we successfully identified four TBPs: *B. (Borreliella) burgdorferi* s.s.; *B. (Borrelia) miyamotoi*; *Anaplasma phagocytophilum*; and *Ehrlichia muris eauclairensis*. These findings were in agreement with 16S data generated using Illumina MiSeq microbiome sequencing. We also demonstrate that NAS successfully enriched for TBP-derived sequences to cover entire bacterial pathogen genomes. Importantly, this approach allowed for clear differentiation between closely related bacterial TBPs (e.g., *B. (Borreliella) burgdorferi* s.s., *B. (Borreliella) mayonii*, and *B. (Borrelia) miyamotoi*), highlighting its potential as a novel approach for metagenomic pathogen surveillance across a variety of arthropod vector species.

## 2. Results

We isolated total genomic DNA from four adult female *I. scapularis* specimens collected at different localities across Minnesota (Figure 1). The V4 region of the bacterial 16S rRNA gene was amplified in all four samples, and microbiome sequencing using the Illumina MiSeq platform revealed that all four ticks were infected with *B. (Borreliella) spp*. Additionally, three of the four ticks exhibited evidence of co-infection with an additional tick-borne agent. In total, 16S Illumina sequencing suggested the likely presence of four bacterial TBPs: *A. phagocytophilum*, *B.* (*Borelliella*) *burgdorferi* s.l., *B.* (*Borrelia*) *miyamotoi*, and *E. muris eauclairensis* across our *I. scapularis* samples (Table 1). The total number of classified 16S reads varied by sample and TBP—ranging from 1,182 to 34,89—suggesting wide variability in the relative abundance of each bacterial pathogen in its respective tick host.

**Table 1.**
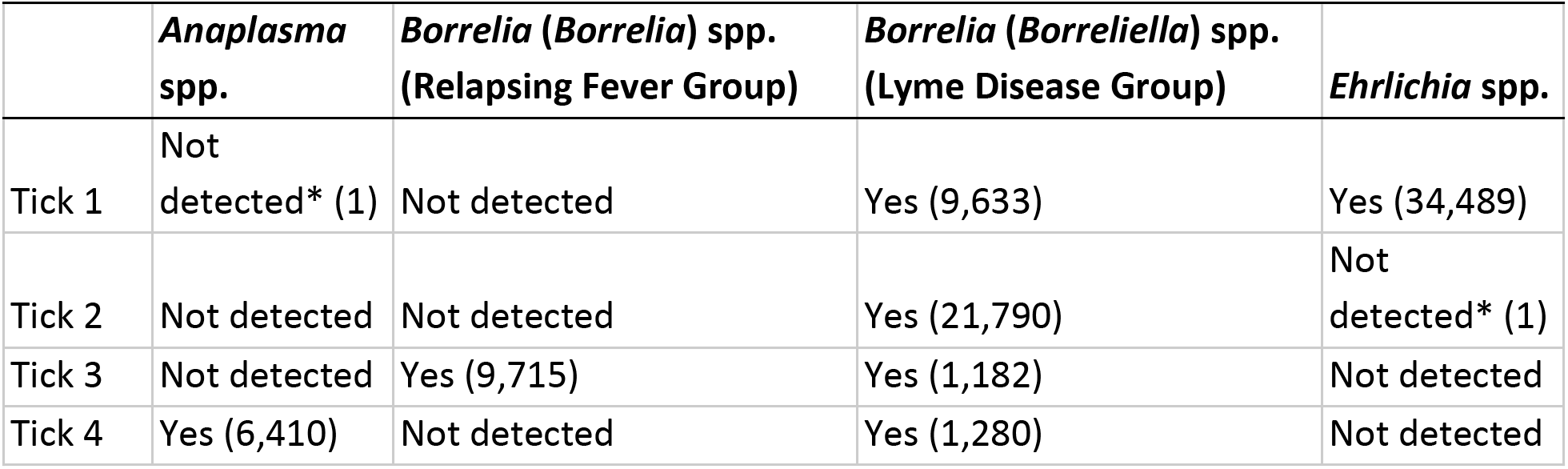
Bacterial tick-borne pathogens detected in adult female *I. scapularis* ticks through 16S microbiome sequencing. Numbers in parentheses indicate the number of assigned 16S Illumina reads. *Taxa for which a single 16S read was assigned are considered likely the result of read misclassification and not indicative of tick infection. See Figure 1 for tick sampling localities in Minnesota, USA.

Next, barcoded nanopore libraries were generated using genomic DNA from the same four *I. scapularis* ticks. To compare the performance of the NAS approach for TBP detection and read enrichment, two identical nanopore libraries were generated. Following pooling of barcoded samples and ligation of nanopore sequencing adapters, both final libraries contained between 140-150 ng of DNA. The libraries were then sequenced on new MinION R9.4 flow cells over a 24 hour runtime. During the sequencing of one library, NAS was enabled. A reference file containing whole genome (and plasmid) sequences of both known and unlikely *Ixodes*-transmitted TBPs was provided for enrichment (Supplemental Table 1). The second nanopore library was sequenced without NAS as a control (Figure 1). The library sequenced with adaptive sampling yielded a total of over 5.5 million reads and 1.9 Gb of sequence data. A comparable amount of total sequencing output was obtained from the control library, with 1.8 Gb of sequencing data generated from a total of 2.9 million reads (Table 2). A notably shorter average read length was obtained from the NAS sequencing run in comparison to the control run (359.0 bp and 627.3 bp, respectively). This difference in average read length was anticipated and likely due to the retention of a significant number of unmapped *I. scapularis* reads in which the initial 200-400 bp were sequenced prior to being rejected from the nanopore.

**Table 2.**
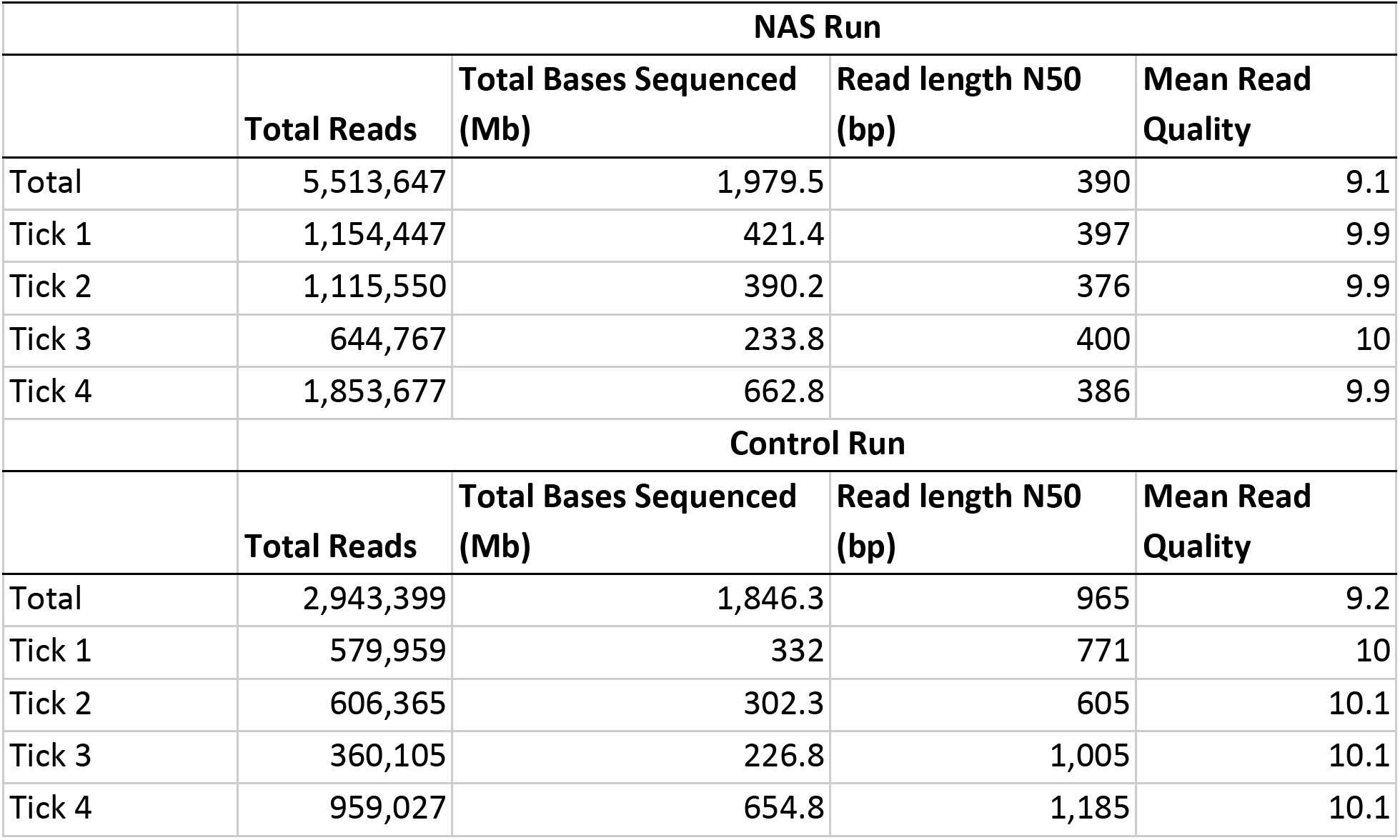
Total sequencing output and output by tick sample generated by adaptive sampling (NAS) and control MinION runs. Reads below quality threshold filtering and unclassified reads without identifiable barcodes are included in total output measurements.

Reads basecalled using the fast basecalling model were aligned to target TBP genomes as sequencing was occurring to provide a real-time assessment of which pathogens were detected. Sequencing of NAS and control libraries yielded a total of 4,216 and 2,631 successfully mapped reads, respectively. The real-time mapping results we obtained were in full agreement with our previous 16S Illumina MiSeq findings in terms of TBPs detected by sample. Importantly, real-time alignment also enabled us to distinguish between the closely-related Lyme Disease Group spirochetes *B.* (*Borreliella*) burgdorferi s.s. and *B.* (*Borreliella*) *mayonii* to confirm that all four ticks were infected with *B.* (*Borreliella*) burgdorferi s.s., a level of taxonomic resolution not attainable from the V4 16S Illumina MiSeq data. Following post-hoc basecalling of raw nanopore reads and the extraction of those reads aligning to our target TBPs, we observed that NAS achieved roughly two-fold total read enrichment over our control sequencing library (Figure 2). This roughly two-fold enrichment was observed to be consistent across all four tick samples and individual TBPs. Similarly, the total number of successfully mapped bases indicated a similar level of sequencing enrichment when NAS was enabled (Supplemental Figure 1). We also noted that numerous reads for both sequencing runs mapped to the genome of the eukaryotic pathogen, *Babesia microti*; however, additional characterization of these reads suggested that they derived from repetitive elements in the *I. scapularis* genome. For this reason, we chose to focus our analyses on characterizing bacterial TBPs only.

**Figure 2.**
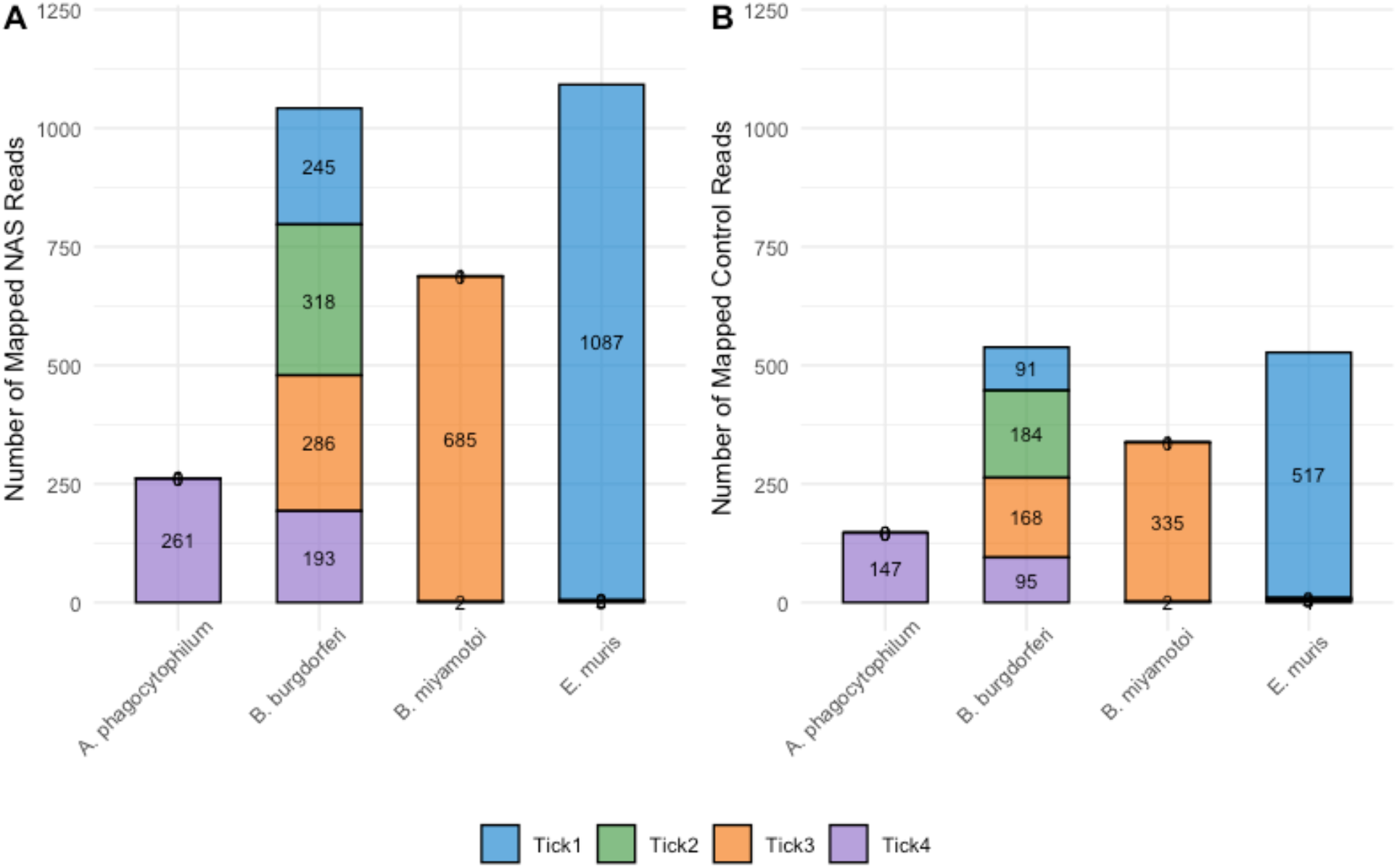
Total nanopore reads mapping to detected bacterial tick-borne pathogen genomes by *I. scapularis* sample (Tick 1 - Tick 4). (A) Number of generated reads through nanopore adaptive sampling (NAS). (B) Successfully mapped reads generated using unenriched control sequencing. Across each of the four bacterial pathogens detected, NAS achieved roughly two-fold read enrichment in comparison to control sequencing.

Total reads were then processed using the software package kraken2, an alternative minimizer-based classification tool independent of our previous mapping and characterization using the minimap2 package for real-time TBP genome alignment [42,43]. This approach for read classification provided an additional and independent assessment of taxonomic diversity found within each *I. scapularis* sample. The output from these analyses supported our previous findings in terms of which bacterial TBPs were detected and the level of sequencing enrichment achieved using NAS (Supplemental Figure 2). Overall, the relative numbers of nanopore reads classified using both minimap2 alignment and the kraken2 pipeline were comparable and consistent between samples and sequencing runs. The kraken2 pipeline was also used to provide additional confirmation of the presence of *B. (Borreliella) burgdorferi* s.s. in our samples. Very few reads were classified as *B. (Borreliella) mayonii*, while >100 reads were classified as *B. (Borreliella) burgdorferi* s.s. for each sample over both NAS and control runs.

After concluding that sequencing via NAS achieved a notable level of TBP sequence enrichment over our control run, we assessed where these reads mapped to corresponding pathogen genomes. We extracted NAS mapping coordinates for all TBP mapped reads and then plotted these across the complete genomes of the corresponding TBP (Figure 3). These visualizations demonstrate that sequencing via NAS resulted in a substantial amount of long-read sequence data with numerous bacterial TBP sequences in excess of 5kb. Additionally, we demonstrate that pathogen-derived NAS sequences spanned the assembled genomes, at varying depths of coverage, of each TBP.

**Figure 3.**
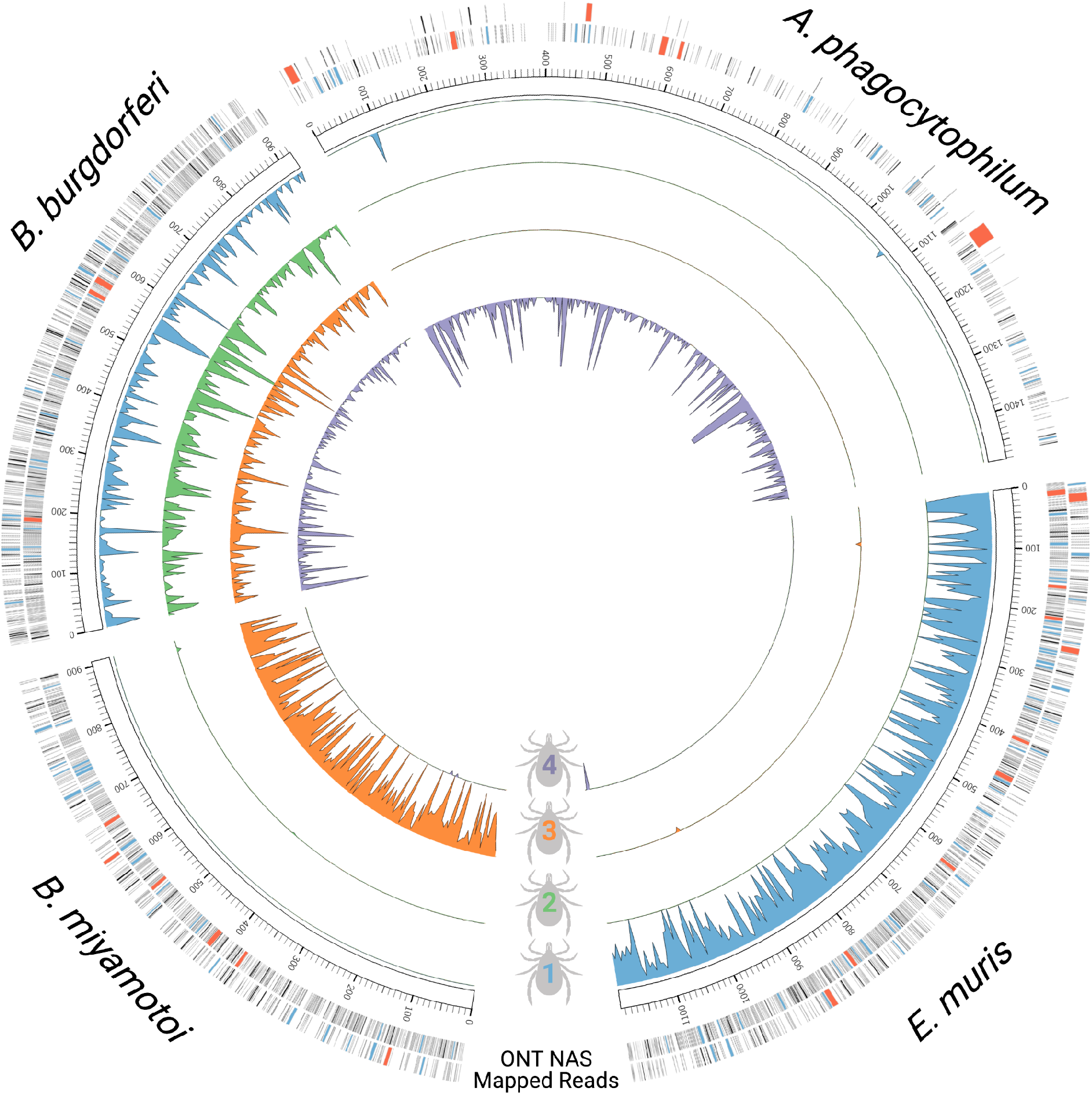
NAS mapping results for four pathogen genomes associated with individual *I. scapularis* ticks. Outermost bands are ONT reads mapping to the genomes of *B. burgdorferi*, *A. phagocytophilum*, *E. muris*, and *B. miyamotoi* with read lengths colored according size (> 5kb indicated in red; > 2kb in blue, and < 2kb in grey). Major tick marks of respective genomes are shown in 100kb segments. Four individual ticks were sampled with NAS and are color-coded in blue (1), green (2), orange (3), and purple (4). Inner bands show mean mapped bases for pathogens associated with each tick sample in a sliding window of 5kb (blue, green, orange, and purple histograms).

## 3. Discussion

Surveillance of vector arthropods for pathogens is an important part of the public health repertoire for combating vector-borne diseases. The U.S. Centers for Disease Control and Prevention have stated that “new tools for preventing tick-borne diseases are urgently needed…” [44]. Nanopore sequencing allows for rapid detection of pathogens with minimal sample preparation and reagent requirements, and can be performed across a variety of field conditions. This facet of the technology brings the possibility of arthropod surveillance sequencing to groups and regions without ready access to laboratory or internet resources. In places with high rates of endemic vector-borne disease, local public health organizations can use nanopore sequencing to rapidly determine the prevalence of pathogens in vector species without needing to rely on the testing facilities of state or national laboratories. Moreover, unlike traditional PCR which allows for pathogen detection by amplifying short regions of template DNA, nanopore sequencing produces long reads that provide substantially more genomic information. As such, nanopore sequencing will see utility in the differentiation of pathogenic and non-pathogenic strains within the same species. For example, human infectious *A. phagocytophilum* strains can be rapidly differentiated from non-human infectious ungulate strains without requiring nested PCR [45]. Genetically similar species, such as spotted fever group rickettsiae (some of which are not human infectious), will be rapidly differentiable in the field rather than relying on PCR amplification followed by sequencing or restriction fragment length polymorphism techniques, as is currently done [46,47].

The recent development of adaptive sampling on ONT sequencing platforms offers even greater potential for sequence-based applications in arthropod vector surveillance. Using NAS, investigators can rapidly deplete arthropod host-associated sequences to enrich metagenomic communities and any vector-borne pathogens contained therein. Additionally, NAS can be used to selectively sequence reads associated with target pathogens which may be substantially less abundant than other microbes. Using ticks as an example, many species contain large quantities of symbiotic bacteria belonging to the same genera as TBPs such as *Rickettsia amblyommatis* in *Amblyomma* spp. or *Francisella* sp. in *Dermacentor* spp. In *I. scapularis*, 10^6^ - 10^7^ *Rickettsia buchneri* may be present in a single female [48] and this symbiont commonly represents >80% of 16S sequences in microbiome studies of female ticks [49–51]. The sheer quantity of genetic material from these bacteria can overwhelm less-abundant or rare sequences in methods used to evaluate the presence of multiple bacterial species within field sampled ticks. Thus, NAS provides a valuable means of selecting against the sequencing of non-target reads in this context.

We leveraged nanopore MinION sequencing in combination with the newly released NAS pipeline to detect and molecularly characterize four TBPs (*B. (Borreliella) burgdorferi s.s.*; *B.* (*Borrelia*) *miyamotoi*; *A. phagocytophilum*; and *E. muris eauclairensis*) associated with four female *I. scapularis* ticks. Using whole genome assemblies for these bacterial pathogens, we confirmed that NAS successfully enriched for each TBP, resulting in an approximately two-fold enrichment of recovered single-molecule sequences over control sequencing (Figure 2; Supplemental Figure 1). Our presence/absence findings were supported by 16S Illumina microbiome sequencing data and an independent method for classifying nanopore reads. Importantly, nanopore sequencing enabled us to achieve a high level of taxonomic resolution in these samples to conclude that all four ticks were infected with *B. (Borreliella) burgdorferi s.s.* and not the closely related species *B. (Borreliella) mayonii.* Although PCR-based tick surveillance approaches can easily provide similar species-level resolution, this approach enabled us to generate large amounts of sequence data spanning entire bacterial TBP genomes. Using NAS, we noted the recovery of extremely long, intact DNA sequences associated with each bacterial TBP. For example, we received a single *A. phagocytophilum* read in excess of 26 kb, in addition to numerous >5 kb reads across all detected TBPs (Figure 3). Such long sequence lengths of individual DNA molecules secured using the NAS bioinformatic pipeline opens the door to genome-scale pathogen surveillance, an opportunity that is not easily explored using traditional methods.

Despite documenting the functionality of NAS for TBP research, we noted that numerous nanopore reads incorrectly aligned against the *Ba. microti* genome, a protozoal eukaryotic pathogen. Additional analysis found that these reads contained repetitive DNA sequences derived from the host *Ixodes* tick. Thus, we recommend caution when interpreting NAS sequence data wherein highly conserved or repetitive sequences have the potential to provide incorrect taxonomic assignment or spurious results. This caveat notwithstanding, nanopore sequencing in combination with adaptive sampling showed considerable promise at detecting bacterial TBPs using unprocessed genomic DNA and limited sample processing steps. As “proof-of-concept”, our study only assessed four field-collected *I. scapularis* ticks; however we anticipate that with optimization of library preparation methods and performance improvements to the NAS pipeline, a substantially greater number of samples may be multiplexed on a single MinION flow cell. As the technologies and computational abilities associated with nanopore sequencing and adaptive sampling continue to improve, a multitude of exciting applications in real-time, metagenomic pathogen biosurveillance are becoming possible over a variety of host-pathogen systems.

## 4. Materials and Methods

### Tick sample collection and nucleic acid extraction

As part of larger sampling efforts, ticks were collected using a standard cloth dragging technique from four collection locations in Minnesota between April 2017 and June 2019. Sampling consisted of dragging a light-colored 1-m^2^ cloth over the ground and low-laying vegetation to attract host-seeking ticks. Ticks attached to the drag cloth were removed using forceps and visually identified to species, developmental stage, and sex. A single adult female *I. scapularis* specimen was selected from each of the four localities sampled to be included in this study. Surface contaminants were removed by sequentially washing ticks in Tween 20 and 0.5% benzalkonium chloride solutions; ticks were next rinsed with distilled water and 70% ethanol, and stored in fresh 70% ethanol prior to nucleic acid extraction. To promote effective tissue lysis, whole ticks were bisected using a sterile 16-gauge needle and incubated in a lysis buffer (i.e., 180 μl Qiagen Buffer ATL and 20 μl proteinase K) at 56°C overnight. The resulting digestion mixture was processed using a DNeasy Blood and Tissue Kit (Qiagen, Hilden, Germany) following manufacturer instructions. DNA extracts were quantified using a Qubit 4 fluorometer (Invitrogen, Carlsbad, United States) and visualized by agarose gel electrophoresis.

### 16S microbiome sequencing

Characterization of microbial diversity was accomplished through amplification of the V4 region of the 16S rRNA gene using the Meta_V4_515F (TCG- TCG - GCA - GCG- TCA- GAT- GTG-TAT-AAG-AGA-CAG-GTG-CCA-GCM-GCC-GCG-GTA-A) and Meta_V4_806R (GTCTCGTGGGCTCGGAGATGTGTATAAGAGACAGGGACTACHVGGGTWTCTAAT) primers.

An initial qPCR step was performed using the Meta_V4_515F and Meta_V4_806R primer pair, and consisted of an initial 95° C denaturation for five minutes, 35 cycles of 98° C denaturation for 20 seconds, 55° C annealing for 15 seconds, and 72° C extension for 60 seconds, and concluded with a final extension for five minutes at 72° C. These data were used to normalize each tick template sample to approximately 167,000 molecules/μl. Next, a conventional PCR using the same primer pair and normalized template was conducted using the following parameters: initial denaturation at 95° C for 5 minutes; 25 cycles of 98° C denaturation for 20 seconds, 55° C annealing for 15 seconds; and 72° C extension for 60 seconds; followed by a 72° C final extension for 5 minutes. A 1:100 dilution of the resulting PCR products was done, and 5 μl of this diluted product was used for a second PCR consisting of forward (AATGATACGGCGACCACCGAGATCTACAC[i5]TCGTCGGCAGCGTC) and reverse (CAAGCAGAAGACGGCATACGAGAT[i7]GTCTCGTGGGCTCGG) Illumina indexing primers for Nextera adapters. The specification for this second PCR were as follows: initial denaturation at 95° C for 5 minutes; 10 cycles of 98° C denaturation for 20 seconds, 55° C annealing for 15 seconds; 72° C extension for 60 seconds; followed by a final 72° C extension for 5 minutes. Second round PCR products were then pooled, denatured with NaOH, diluted to 8 pM using Illumina HT1 buffer, spiked with 15% PhiX, and denatured at 96° C for 2 minutes. The prepared Illumina library was sequenced on the Illumina MiSeq system with a 600 cycle v3 kit.

### Nanopore library preparation

Genomic DNA libraries were generated following the Sequencing Ligation Kit SQK-LSK109 (ONT) protocol, using between 0.8 and 1.1 μg of total input DNA from each of the four tick samples. Repair and end-prep of DNA molecules was carried out using NEBNext FFPE DNA Repair and Ultra II End Repair/dA-Tailing modules (New England Biolabs Inc., Ipswich, United States), incubated at 20° C for 5 minutes followed by heat-inactivation at 65° C for an additional 5 minutes. Samples were purified using AMPure XP beads (Beckman Coulter, Indianapolis, United States) on a magnetic separation rack at a ratio of 1.8:1 beads-to-sample. Barcodes were ligated to DNAs for each of the four tick samples using Blunt/TA Ligase Master Mix (New England Biolabs Inc., Ipswich, United States) and the EXP-NBD104 Native Barcoding Expansion Kit (ONT). Following barcode ligation, samples were quantified on a Qubit 4 fluorometer (Invitrogen, Carlsbad, United States) and equimolar amounts of each barcoded sample were pooled and used to generate two identical nanopore libraries. Sequencing adapters were ligated to the libraries using NEBNext Quick Ligation Reaction Buffer/Quick T4 DNA Ligase (New England Biolabs Inc., Ipswich, United States); ONT Short Fragment Buffer was used during the final library wash to retain fragments of all lengths. Final pooled libraries were eluted in a volume of 15 μl at 37° C, quantified as above, and stored at 4° C prior to sequencing.

### MinION sequencing and basecalling

Both pooled libraries were sequenced over two separate experiments, with each experiment employing a new FLO-MIN106 flow cell using R9.4 sequencing chemistry (ONT). Each flow cell was checked immediately prior to sequencing to ensure that >1,200 active pores were available for sequencing. Flow cells were primed and loaded on an ONT MinION Mk1B device connected to a custom desktop computer operating on Ubuntu 18.04 LTS and with the following hardware specifications: a GeForce 2080Ti GPU (Nvidia, Santa Clara, United States); a 24-core Ryzen 3900x CPU (Advanced Micro Devices, Inc., Santa Clara, United States); 64 Gb RAM; a 1 TB SSD; and 15 TB total onboard storage. Sequencing experiments were initiated and monitored using MinKNOW software, v20.10.3. Adaptive sampling to enrich for pathogen-associated reads was carried out for one library using ReadUntil (https://github.com/nanoporetech/read_until_api), integrated into the MinKNOW platform. To achieve adaptive sampling, reference files in .mmi and .bed format containing whole genome sequences and coordinates of potential *I. scapularis*-transmitted pathogens were supplied for real-time enrichment; associated plasmid sequences were also included. These pathogens consisted of *B. (Borreliella) burgdorferi* s.s.; *B. (Borreliella) mayonii*; *B. (Borrelia) miyamotoi*; *A. phagocytophilum*; *E. muris eauclairensis; Ba. microti;* Powassan virus, as well as the unlikely *I. scapularis*-transmitted agents *Bartonella henselae* and the mitogenome for *Dirofilaria immitis* (Supplemental Table 2). As a control, the second library was sequenced without adaptive sampling enrichment. For both MinION libraries, real-time alignment using the same NAS reference file was enabled in MinKNOW to allow mapping of reads against our TBP genomes using the minimap2 software package [43]. Reads mapped during sequencing in real-time were basecalled using the fast basecalling model in Guppy; however post-hoc high accuracy basecalling was performed prior to all subsequent analyses. Both NAS and control sequencing runs were allowed to proceed for 24 hours to allow each library to be sequenced to near completion. After sequencing, raw reads in Fast5 format were re-basecalled and demultiplexed using Guppy, v4.2.2, in GPU mode using the high accuracy (HAC) basecalling model. Raw nanopore sequence data from both NAS and control sequencing runs are deposited in the NCBI Short Read Archive (SRA), under project number [Accession number pending publication].

### Bioinformatic processing

Paired-end raw 16S reads generated on the Illumina MiSeq platform were demultiplexed and Nextera adapters were trimmed. The resulting reads were processed using the DADA2 pipeline (https://github.com/benjjneb/dada2) to taxonomically assign 16S amplicons to the level of the bacterial genus. The resulting amplicon sequence variant file was queried for pathogenic agents of interest (e.g., *Anaplasma*, *Borrelia*, *Borreliella*, *Ehrlichia*) to determine pathogen presence and the relative abundance of raw V4 16S reads for each sample. Quality statistics from both NAS and control nanopore MinION sequencing runs were assessed using NanoPlot (https://github.com/wdecoster/NanoPlot) and NanoComp (https://github.com/wdecoster/nanocomp) software packages [52]. For both sequencing runs, demultiplexed and HAC basecalled FASTQs were concatenated by sample, and passed reads with a mean read quality score of greater than 7 were retained for downstream analyses. Adaptive sampling .csv and sequencing summary .txt output files were queried to assess which nanopore reads from both sequencing runs successfully aligned to our target TBP genomes. Independent taxonomic classification was performed using the kraken2 software package, v2.1.1, (https://github.com/DerrickWood/kraken2) [42] and the pre-assembled “PlusPF” database (https://benlangmead.github.io/aws-indexes/k2) for assessing archaeal, bacterial, viral, protozoal, and fungal metagenomic diversity. For each tick sample, passed nanopore reads in FASTQ format were analyzed against the kraken2 PlusPF database, and kraken2 output reports were queried to determine the number of classified reads belonging to TBP taxa of interest.

## 5. Conclusions

Using nanopore MinION sequencing, in combination with the recently developed NAS bioinformatic pipeline, we selectively sequenced bacterial TBP DNA sequences from infected *I. scapularis* ticks. Across the four individual ticks assessed, we detected a total of four bacterial agents: *Borrelia* (*Borreliella*) *burgdorferi* s.s.; *Borrelia* (*Borrelia*) *miyamotoi*; *Anaplasma phagocytophilum*; and *Ehrlichia muris eauclairensis*. The presence of these TBPs was confirmed with 16S metagenomic sequencing using Illumina MiSeq. In comparison to our control sequencing library, adaptive sampling achieved roughly two-fold sequence enrichment for our target TBPs and was able to provide clear differentiation between closely related bacterial TBPs. Independent taxonomic classification of our nanopore reads confirmed our pathogen presence findings and demonstrated similar levels of target read enrichment using the NAS pipeline. Importantly, we showed that NAS generated individual nanopore reads thousands of bases in length that collectively spanned the entire genomes of our TBPs of interest. We posit that current and future iterations of NAS will facilitate a wide variety of real-time metagenomic biosurveillance initiatives.

## Author Contributions

Conceptualization: PAL, JDO, EJK

Investigation: EJK, LLL, CF, BK, JDO, PAL

Writing - original draft preparation: EJK, PAL, BK, JDO

Writing - reviewing and editing: EJK, LLL, CF, BK, JDO, PAL

All authors have read and agreed to the published version of the manuscript.

## Data Availability Statement

Raw data will be deposited on NCBI SRA prior to publication and is available to reviewers upon request

## Acknowledgements

We thank Suzanne Stone for logistical support of the research performed herein. The Minnesota Supercomputing Institute (MSI) at the University of Minnesota provided key resources to support bioinformatic analyses and data storage. The University of Minnesota Genomic Center (UMGC) performed Illumina library preparation and 16S sequencing. We thank Trevor Gould of the UMGC for bioinformatic analyses of 16S data.

