## Supplemental Table 1 for "Enabling metagenomic surveillance for bacterial tick-borne pathogens using nanopore sequencing with adaptive sampling"

**Supplemental Table 1.** Whole genome and plasmid sequences of known and potential tick-borne pathogens included in enrichment FASTA file for adaptive sampling enrichment.

| Sequence Number | Organism | Group | Strain | Sequence Type | Accession Number | Length (bp) |
|---|---|---|---|---|---|---|
| 1 | *B. burgdorferi* s.s. | Bacteria | JD1 | Complete Genome | CP002312.1 | 922,801 |
| 2 | *B. burgdorferi* s.s. | Bacteria | JD1 | Plasmid | CP002316.1 | 26,525 |
| 3 | *B. burgdorferi* s.s. | Bacteria | JD1 | Plasmid | CP002310.1 | 60,739 |
| 4 | *B. burgdorferi* s.s. | Bacteria | JD1 | Plasmid | CP002323.1 | 30,314 |
| 5 | *B. burgdorferi* s.s. | Bacteria | JD1 | Plasmid | CP002309.1 | 29,902 |
| 6 | *B. burgdorferi* s.s. | Bacteria | JD1 | Plasmid | CP002321.1 | 30,019 |
| 7 | *B. burgdorferi* s.s. | Bacteria | JD1 | Plasmid | CP002320.1 | 29,707 |
| 8 | *B. burgdorferi* s.s. | Bacteria | JD1 | Plasmid | CP002307.1 | 30,804 |
| 9 | *B. burgdorferi* s.s. | Bacteria | JD1 | Plasmid | CP002311.1 | 31,085 |
| 10 | *B. burgdorferi* s.s. | Bacteria | JD1 | Plasmid | CP002322.1 | 30,697 |
| 11 | *B. burgdorferi* s.s. | Bacteria | JD1 | Plasmid | CP002313.1 | 17,522 |
| 12 | *B. burgdorferi* s.s. | Bacteria | JD1 | Plasmid | CP002308.1 | 22,827 |
| 13 | *B. burgdorferi* s.s. | Bacteria | JD1 | Plasmid | CP002306.1 | 24,354 |
| 14 | *B. burgdorferi* s.s. | Bacteria | JD1 | Plasmid | CP002314.1 | 28,797 |
| 15 | *B. burgdorferi* s.s. | Bacteria | JD1 | Plasmid | CP002318.1 | 30,096 |
| 16 | *B. burgdorferi* s.s. | Bacteria | JD1 | Plasmid | CP002317.1 | 24,614 |
| 17 | *B. burgdorferi* s.s. | Bacteria | JD1 | Plasmid | CP002324.1 | 26,981 |
| 18 | *B. burgdorferi* s.s. | Bacteria | JD1 | Plasmid | CP002319.1 | 29,993 |
| 19 | *B. burgdorferi* s.s. | Bacteria | JD1 | Plasmid | CP002315.1 | 22,808 |
| 20 | *B. burgdorferi* s.s. | Bacteria | JD1 | Plasmid | CP002325.1 | 27,590 |
| 21 | *B. burgdorferi* s.s. | Bacteria | JD1 | Plasmid | CP001652.1 | 52,916 |
| 22 | *A. phagocytophilum* | Bacteria | JM | Complete Genome | NC_021880.1 | 1,481,598 |
| 23 | *Ba. microti* | Protozoa | RI | Chromosome 1 | FO082871.1 | 1,304,281 |
| 24 | *Ba. microti* | Protozoa | RI | Chromosome 2 | FO082872.1 | 1,508,385 |
| 25 | *Ba. microti* | Protozoa | RI | Chromosome 3 | LN871598.1 | 1,766,409 |
| 26 | *Ba. microti* | Protozoa | RI | Chromosome 4 | LN871599.1 | 1,816,206 |
| 27 | *Ba. microti* | Protozoa | RI | Mitochondrial Genome 1 | LN871602.1 | 10,547 |
| 28 | *Ba. microti* | Protozoa | RI | Mitochondrial Genome 2 | LN871600.2 | 11,149 |
| 29 | *Ba. microti* | Protozoa | RI | Mitochondrial Genome 3 | LN871601.1 | 10,547 |
| 30 | *Ba. microti* | Protozoa | RI | Mitochondrial Genome 4 | LN871603.1 | 10,547 |
| 31 | *Ba. microti* | Protozoa | RI | Apicoplast Genome | LK028575.1 | 28,657 |
| 32 | *E. muris* | Bacteria | AS145 | Complete Genome | CP006917.1 | 1,196,717 |
| 33 | *B. miyamotoi* | Bacteria | LB-2001 | Complete Genome | CP006647.2 | 907,293 |
| 34 | *B. miyamotoi* | Bacteria | LB-2001 | Plasmid | CP010328.2 | 30,673 |
| 35 | *B. mayonii* | Bacteria | MN14-15339 | Complete Genome | CP015796.1 | 904,387 |

| | | | | | | |
|---|---|---|---|---|---|---|
| 36 | *B. mayonii* | Bacteria | MN14-15339 | Plasmid | CP015797.1 | 26,835 |
| 37 | *B. mayonii* | Bacteria | MN14-15339 | Plasmid | CP015798.1 | 30,433 |
| 38 | *B. mayonii* | Bacteria | MN14-15339 | Plasmid | CP015799.1 | 27,866 |
| 39 | *B. mayonii* | Bacteria | MN14-15339 | Plasmid | CP015800.1 | 30,330 |
| 40 | *B. mayonii* | Bacteria | MN14-15339 | Plasmid | CP015801.1 | 30,406 |
| 41 | *B. mayonii* | Bacteria | MN14-15339 | Plasmid | CP015802.1 | 17,017 |
| 42 | *B. mayonii* | Bacteria | MN14-15339 | Plasmid | CP015962.1 | 8,307 |
| 43 | *B. mayonii* | Bacteria | MN14-15339 | Plasmid | CP015803.1 | 23,759 |
| 44 | *B. mayonii* | Bacteria | MN14-15339 | Plasmid | CP015804.1 | 24,696 |
| 45 | *B. mayonii* | Bacteria | MN14-15339 | Plasmid | CP015805.1 | 19,566 |
| 46 | *B. mayonii* | Bacteria | MN14-15339 | Plasmid | CP015806.1 | 27,809 |
| 47 | *B. mayonii* | Bacteria | MN14-15339 | Plasmid | CP015807.1 | 32,269 |
| 48 | *B. mayonii* | Bacteria | MN14-15339 | Plasmid | CP015808.1 | 45,035 |
| 49 | *B. mayonii* | Bacteria | MN14-15339 | Plasmid | CP015809.1 | 53,343 |
| 50 | Powassan virus | Virus | ctb30 | Complete Genome | AF311056.1 | 10,800 |
| 51 | *Dirofilaria immitis* | Helminth | N/A | Mitochondrial Genome | NC_005305.1 | 13,814 |
| 52 | *Bartonella henselae* | Bacteria | MVT02 | Complete Genome | LN879429.1 | 1,905,383 |
