## Supplemental Figure 1 for "Enabling metagenomic surveillance for bacterial tick-borne pathogens using nanopore sequencing with adaptive sampling"

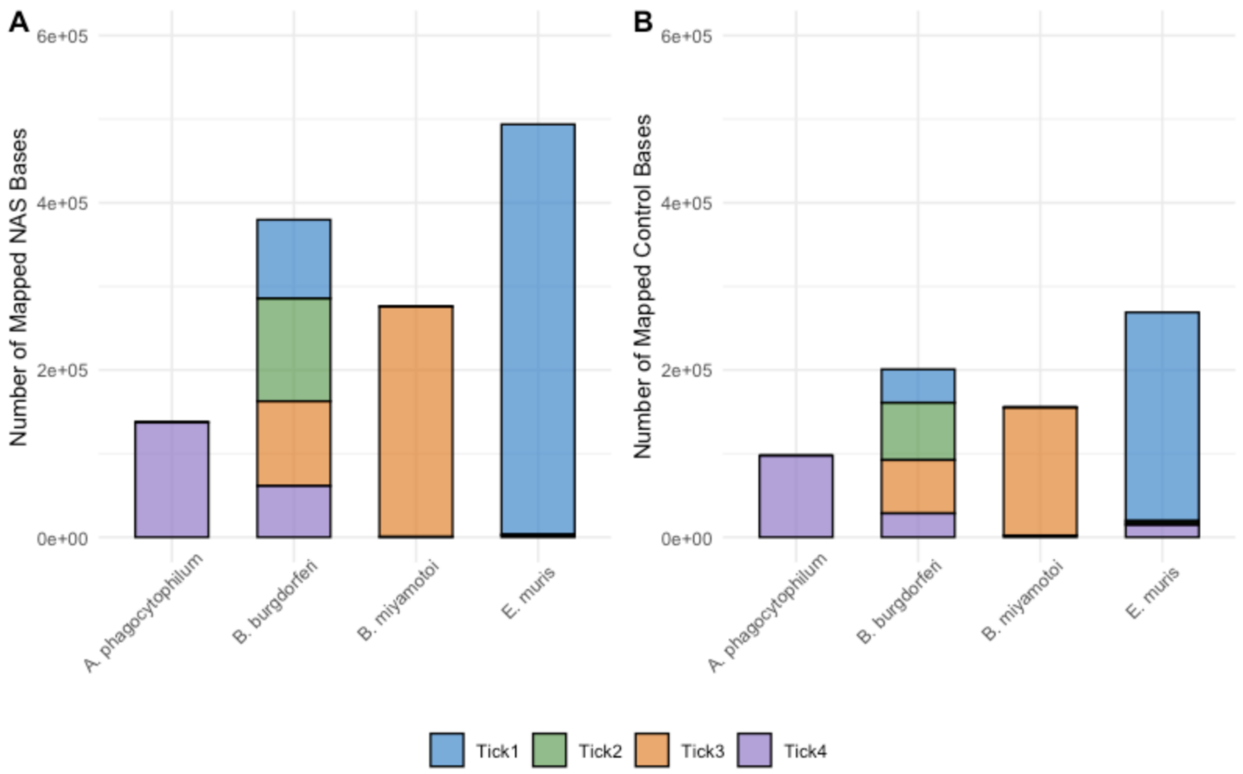


**Supplemental Figure 1.** Total number of nanopore bases mapping to each bacterial tick-borne pathogen genome for each tick sample. (A) Number of mapped bases generated during NAS sequencing run. (B) Mapped bases generated during unenriched control sequencing run.
