## Supplemental Figure 2 for "Enabling metagenomic surveillance for bacterial tick-borne pathogens using nanopore sequencing with adaptive sampling"

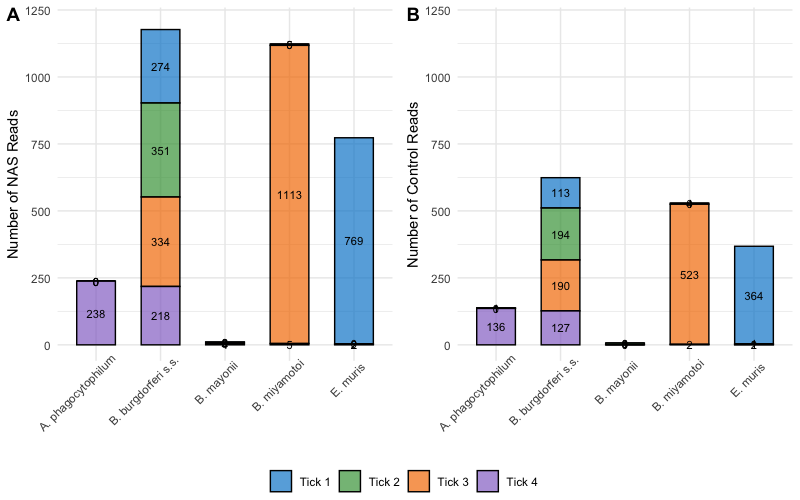

**Supplemental Figure 2.** Number of nanopore reads assigned taxonomic classification at the level of pathogen species using the kraken2 pipeline. The tick-borne agent *B.* (*Borreliella*) *mayonii* was not detected in these samples and was included to confirm of sample infections with the related pathogen *B. (Borreliella) burgdorferi* s.s. (A) Number of classified NAS reads (B) Number of classified control reads.
